# Divergent Golgi Trafficking Limits B cell-Mediated IgG Sialylation

**DOI:** 10.1101/2022.01.25.477787

**Authors:** Leandre M. Glendenning, Julie Y. Zhou, Kalob M. Reynero, Brian A. Cobb

## Abstract

The degree of α2,6-linked sialylation on IgG glycans is associated with a variety of inflammatory conditions and is thought to drive IgG anti-inflammatory activity. Previous findings revealed that ablation of β-galactoside α2,6-sialyltransferase 1 (ST6Gal1) in B cells failed to alter IgG sialylation *in vivo,* yet resulted in the loss of B cell surface α2,6 sialylation, suggesting divergent pathways for IgG and cell surface glycoprotein glycosylation and trafficking. Employing both B cell hybridomas and *ex vivo* murine B cells, we discovered that IgG was poorly sialylated by ST6Gal1 and highly core fucosylated by α1,6-fucosyltransferase 8 (Fut8) in cell culture. In contrast, IgG-producing cells showed the opposite pattern by flow cytometry, with high cell surface α2,6 sialylation and low α1,6 fucosylation. Paired studies further revealed that *ex vivo* B cell-produced IgG carried significantly less sialylation compared to IgG isolated from the plasma of matched animals, providing evidence that sialylation increases after release *in vivo.* Finally, confocal analyses demonstrated that IgG poorly localized to subcellular compartments rich in sialylation and ST6Gal1, and strongly to regions rich in fucosylation and Fut8. These findings support a model in which IgG subcellular trafficking diverges from the canonical secretory pathway by promoting Fut8-mediated core fucosylation and limiting exposure to and modification by ST6Gal1, providing a mechanism for why B cell-expressed ST6Gal1 is dispensable for IgG sialylation *in vivo.*

## Introduction

IgG sialylation has been implicated in a variety of human conditions, including infectious diseases and autoimmunity (reviewed in [1]). Human IgG contains a single conserved N-glycosylation site at Asn297 on the Fc domain, and the associated N-glycans are almost always biantennary, with varying amounts of bisecting GlcNAc, galactosylation, core α1,6 fucosylation, and terminal α2,6-linked sialylation [2]. The complexity and branching of this glycan impacts the pro- or anti-inflammatory effector functions of the molecule, likely through altering the ability of the molecule to bind to inhibitory and activating Fcγ receptors [3]. Early studies examining IgG glycosylation in the context of rheumatoid arthritis (RA) found that loss of galactosylation and sialylation were associated with disease [4], and other studies suggest that the addition of sialic acid converts IgG to an anti-inflammatory effector glycoprotein [5]. Further studies have linked a lack of sialylation and galactosylation to a variety of other diseases, including as a biomarker for adverse cardiac events in HIV^+^ patients [6].

Literature describing a consistent association between various inflammatory diseases and changes in IgG sialylation suggests that the addition of sialic acid is a process that is tightly regulated, with widespread immunological downstream effects. Studies involving patients with rheumatoid arthritis document dynamic changes in IgG glycosylation, as RA goes into remission during pregnancy, concomitant with an increase in sialylation on the Fc region. Following childbirth, RA symptoms return, along with a decrease in IgG sialylation and galactosylation [7]. This suggests that glycosylation of IgG and other proteins can be dynamically modified, reflecting a tightly regulated homeostatic role for glycosylation in the context of inflammation. Although associations between sialylation and various diseases have been extensively studied, the mechanisms behind the regulation of sialylation remains poorly understood. Currently, regulation of sialylation is studied primarily in the context of expression of the relevant enzymes [8, 9]; however, further molecular detail on the mechanism of regulation is lacking.

We previously created a B cell-specific conditional knockout (BcKO) of the sialyltransferase ST6Gal1 in mice [10], which is responsible for the addition of terminal sialic acid in an α2,6 linkage. These mice lack ST6Gal1 throughout the B cell compartment, as shown through a dramatic decrease in B cell surface sialylation and concomitant increase in terminal galactose. Although expression of ST6Gal1 in the B cell was necessary for surface glycoprotein sialylation and likely CD22 clustering and B cell receptor signaling regulation, it was not necessary for IgG sialylation. In fact, the IgG localized to the plasma of these mice were normally sialylated [10]. These data suggested not only that sialic acids are added to IgG following secretion, but also that the B cells are potentially unable to sialylate IgG themselves.

The canonical understanding of the secretory pathway indicates that glycoproteins are sialylated in the fr*ons*-Golgi network as part of a stepwise glycan maturation process. Given the divergence of plasma-localized IgG and B cell surface glycosylation in WT and BcKO mice in terms of α2,6 sialylation, we hypothesized that IgG normally lacks intracellular exposure to ST6Gal1, thereby limiting B cell-mediated sialylation. To explore this hypothesis, we employed three mouse hybridoma strains which produce mouse IgG1, IgG2a, or IgG3, as well as primary cells from WT mice. We examined the glycosylation status of IgG and other cellular proteins and identified significant discrepancies. IgG was characterized by low α2,6 sialylation and high α1,6 core fucosylation, yet B cell surface proteins were the opposite, with high sialylation and low core fucosylation. Confocal microscopy further revealed that IgG strongly localized to intracellular compartments rich in core fucose, but poorly to compartments rich in both α2,6 sialylation and the ST6Gal1 enzyme. These data support a model in which IgG glycosylation is driven by intracellular trafficking in B cells, thereby explaining why B cell-expressed ST6Gal1 is dispensable for IgG sialylation *in vivo.*

## Materials & Methods

### Cells and General Culture Conditions

Hybridoma cell lines were obtained from ATCC and chosen based on the IgG subclass produced. These were KL295 (Clone CRL-1996; mouse IgG1), HB55 (Clone L243; mouse IgG2a), and HKPEG1 (Clone CCL-189; mouse IgG3). All cell lines were cultured in Advanced DMEM (Gibco) supplemented with 10 % fetal bovine serum, β-mercaptoethanol, L-glutamine, and penicillin/streptomycin. Primary B cells were obtained by harvesting mouse spleens, which were mashed through a 100 μm filter and sorted by CD19 positive selection using magnetic beads (Miltenyi). Following separation, primary B cells were cultured in Advanced RPMI (Gibco) supplemented with 5 % fetal bovine serum, β-mercaptoethanol, and penicillin/streptomycin.

### Flow cytometry

Flow cytometry was performed on cells isolated from culture by spinning down at 300 x g for 5 minutes. Cells were washed three times with PBS, filtered through a 70 μm nylon filter, and blocked for 30 minutes in Carbohydrate Free Blocking Solution (Vector Labs). Cells were stained with the following fluorescein-conjugated lectins (all Vector Labs): SNA, ConA, PHA-E, PHA-L, LCA, UEA-I, MAL-I, and ECL. Flow cytometry was run on an Attune NxT (Case Comprehensive Cancer Center Cytometry & Microscopy Shared Resource; NIH grant S10-NIH OD021559) and data were analyzed using FlowJo.

### B cell stimulation

Mice were sacrificed by CO2 inhalation according to a CWRU-approved IACUC protocol, and blood was obtained via cardiac blood draw into 3.2% sodium citrate to prevent coagulation. Plasma was generated from blood by taking the liquid fraction following centrifugation at 2500 x g for 15 minutes. B cells were obtained as described and plated in round-bottom 96-well plates at a density of 100,000 cells/well. The cells were stimulated by the addition of 100 ng/mL anti-CD40 (Clone 1C10; eBioscience) and 1 ng/mL *E. coli* LPS (Invivogen) to the media, and incubated for 7 days prior to media harvest.

### IgG purification

IgG was purified from cell culture media or plasma samples by separation over a HiTrap Protein A HP Antibody Purification Column (GE Life Sciences) fitted to a GE Life Sciences Akta Purifier 10 HPLC. Binding and washing were done with 10 mM Tris, pH 7.5, and 150 mM NaCl. Elution of IgG was done with 50 mM Citrate, pH 4.5, and 150 mM NaCl. Purified IgG was buffered-exchanged into PBS (ELISA and long-term storage) or water (for HPLC glycan analyses), and purity validated by SDS-PAGE.

### Glycan HPLC analysis

IgG monosaccharide composition was performed in two steps. First, sialic acids were quantified by hydrolyzing sialic acids in 2 M acetic acid at 80 °C for 3 hours. Samples were dried under nitrogen gas, resuspended in HPLC-grade water, sonicated briefly, and analyzed by high performance anion exchange chromatography (PA-20 column; Dionex) with pulsed amperometric detection (HPAEC-PAD) on a Dionex ICS-5000 HPLC System with a linear gradient of 1 M sodium acetate in 100 mM NaOH from 7 % to 30 %. Second, neutral monosaccharide composition was quantified by hydrolyzing glycans in 2 N trifluoroacetic acid at 100 °C for 4 hours. Samples were dried under nitrogen gas, resuspended in HPLC-grade water, sonicated briefly, and analyzed by HPAEC-PAD with isocratic elution in 10 mM NaOH.

Hydrophilic-interaction liquid chromatography (HILIC) analysis of intact N-glycans was performed using 2-aminobenzamide (2-AB) labeling and separation on a Zorbax NH2 column (Agilent), essentially as described elsewhere [11, 12], glycoproteins were digested with trypsin (ThermoFisher) overnight at 37°C. Trypsin was inactivated by heating to 95 °C for 10 min. N-glycans were enzymatically removed from the underlying peptides with PNGase F (NEB) overnight at 37 °C. Glycans were separated from peptides by reversed-phase chromatography through a C18 cartridge (ThermoFisher) and dried in a lyophilizer (LabConco) overnight. Purified glycan was labeled using 50 μg/μL 2-AB and 60 μg/μL sodium cyanoborohydride in a 3:7 solution of acetic acid and DMSO at 65°C for 3 h. Unreacted label was removed on a G-10 desalting column (GE Healthcare). HILIC analysis was performed using a 36 % to 45 % gradient of 100 mM ammonium formate, pH 4.4 and acetonitrile. 2-AB fluorescence was detected with an excitation of 330 nm and emission of 420 nm.

### Lectin ELISA

Lectin ELISA was performed on purified IgG, as has been published with plasma samples [6]. Briefly, purified IgG was diluted to 1 mg/mL in carbonate coating buffer (100 mM NaHCO_3_, 30 mM NaCO_3_, pH 9.5), pipetted into a 96-well high-binding ELISA plate (Microlon High Binding; Greiner BioOne), and incubated overnight at 4 °C. The plate was blocked with carbohydrate free blocking solution (Vector Labs) for 1 hour at room temperature. Biotinylated lectins (Vector Labs) were diluted to 1 μg/mL in carbohydrate free blocking solution and incubated on the plate for 1 hour at room temperature. Signal was detected using europium-conjugated streptavidin (Perkin Elmer) and time-resolved fluorescence as measured in a Victor V3 1420 multilabel plate reader.

### Expression level of enzymes

RNA transcripts were isolated from cells using a Qiagen RNEasy Mini RNA isolation kit (Qiagen) per the manufacturer’s instructions. RNA was then converted to cDNA using a RevertAid First Strand cDNA synthesis kit (ThermoFisher). 1.5 μg of cDNA was then plated for quantitative PCR with Taqman Fast Advanced Master Mix (ThermoFisher). The primer used against mouse ST6Gal1 was Taqman Mm00486119_m1 and expression was normalized against β2-microglobulin expression with primer probe Mm00437762_m1 (ThermoFisher).

### Analysis of whole cell extracts

Cells were isolated from media by centrifugation at 300 x g for 5 minutes. They were flash-frozen in liquid nitrogen, and stored at −80 to preserve protein integrity. They were then thawed and resuspended in 2 mL of H2O to provide osmotic pressure. Using a Dounce homogenizer, cells were mechanically lysed and centrifuged for 15 min. at 10,000 x g to remove all cellular debris. Sialylation status of whole cell protein extract was then analyzed by lectin ELISA as described.

### Confocal imaging and co-localization analyses

Cells were isolated from media by spinning at 300 x g for 5 minutes. They were washed three times with PBS, then fixed by incubating in 4 % paraformaldehyde for 30 minutes at 4 °C. Cells were washed with PBS, then permeabilized by incubating in 0.1 % Triton-X-100 in PBS for 30 minutes at 4 °C. Next, cells were stained for 3 hours with Phalloidin-iFluor 488 Reagent (Abcam), anti-ST6Gal1 antibody (R&D Systems) labelled with AF647 using a ThermoFisher Protein Labelling Kit (ThermoFisher) and fluorescein-conjugated lectins (all Vector Labs): SNA, LCA, PHA-L, or ConA. Slides were mounted using VECTASHIELD Hardset Antifade Mounting Medium (Vector Labs). Confocal imaging was performed on a SP5 Laser Scanning Confocal Microscope (Leica), and colocalization analyses were performed using ImageJ’s Coloc2 package.

### Data and statistical analysis

All data were analyzed and plotted using GraphPad Prism. Statistical analyses used were Student’s T-test or one-way ANOVA where appropriate (\**P* < .05; \*\**P* < .01; \*\*\**P* < .001; \*\*\*\**P* < 0.0001).

## Results

### Hybridoma-Produced IgG is Poorly Sialylated

In order to understand the relationship between B cell expression of IgG and compositional modulation of its associated N-linked glycans, we chose three murine B cell hybridoma cell lines producing IgG1, IgG2a, or IgG3 and examined their glycosylation using a variety of techniques. IgG was purified from culture supernatants by Protein A chromatography and purity was validated by SDS-PAGE (not shown). Intact IgG glycans were released and labeled with 2-AB and analyzed by HILIC in comparison to human IgG glycans produced and labeled at the National Center for Functional Glycomics. We found that the N-glycans from all three IgG subclasses were very low in sialic acid compared to the human IgG isolated from donor plasma (Figs 1A–1B). Monosaccharide compositional analysis by HPAEC-PAD further demonstrated the proportional molar ratio of sugars reflective of a highly fucosylated and galactosylated bi-antennary N-glycan that lacks significant sialylation (Fig 1C). In fact, sialic acid was below the limit of detection in this assay.

**Figure 1.**
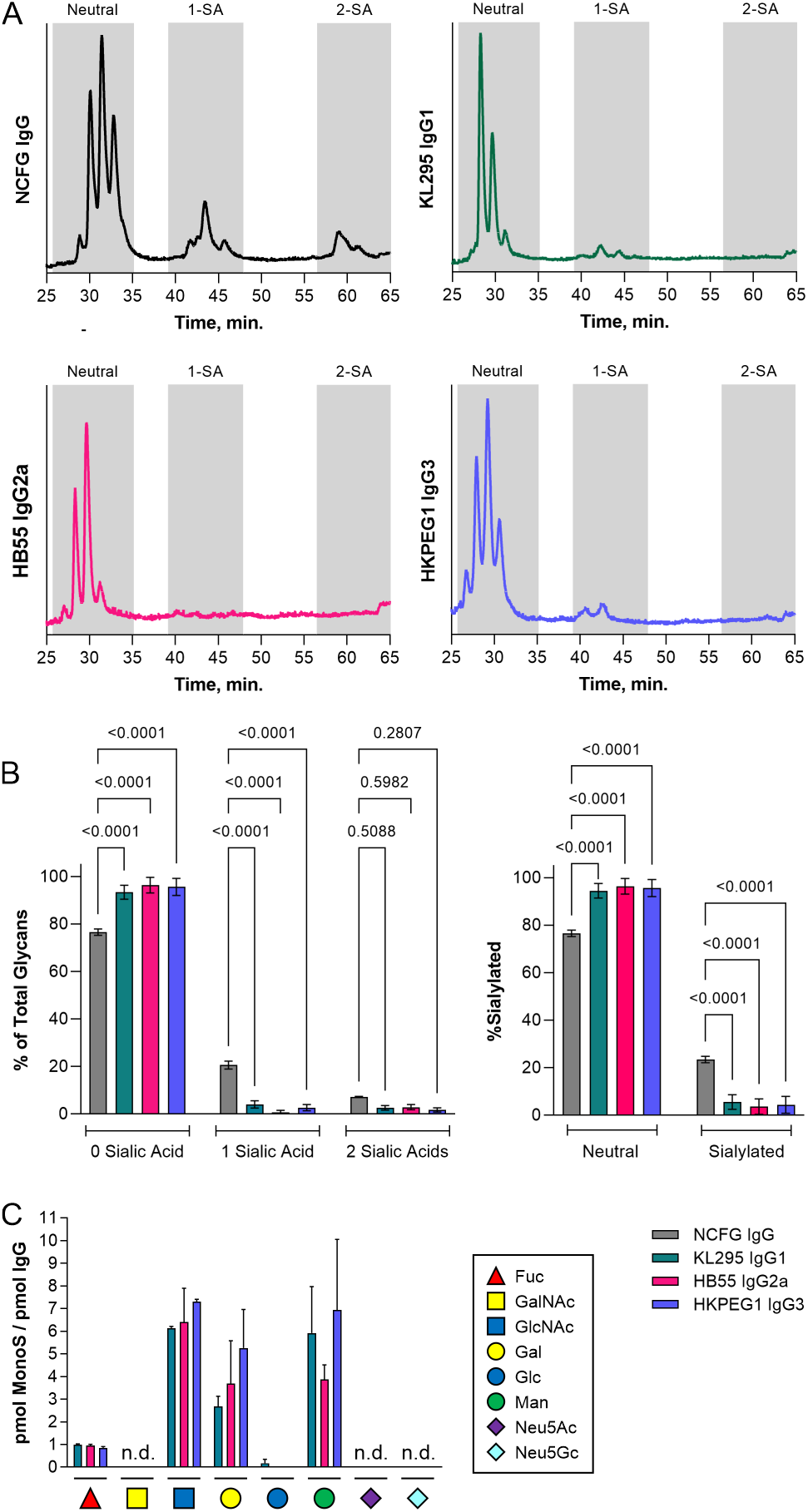
Hybridoma IgG lacks significant sialylation. (A) Representative HILIC HPLC elution profiles of N-glycans from serum-purified human IgG compared to IgG1, IgG2a, and IgG3 produced from KL295, HB55, and HKPEG1 murine hybridomas respectively. Shaded areas indicate oligosaccharides with zero, one, or two sialic acids. (B) Replicate data from HILIC analysis from each IgG sample graphed either as a function of the number of sialic acids (left) or neutral versus total sialylated species (right). N = 5 for the human IgG controls; N = 9 for hybridoma IgGs. (C) Monosaccharide compositional analysis by HPAEC-PAD. Sample marked n.d. were below the limit of detection. N = 3 for neutral monosaccharides. N = 6 for sialic acids. All error bars show SEM.

Using our previously benchmarked lectin-based ELISA-styled assay for the assessment of glycans on glycoproteins [6], we next measured IgG binding by ConA (mannose), LCA (α1,6 core fucose), PHA-E (bisecting GlcNAc), PHA-L (β1,6 GlcNAc branching), and SNA (α2,6 sialylation) at various concentrations in comparison to the heavily glycosylated glycoprotein bovine fetuin as a control (Fig 2A). Using the 100 ng IgG data across all lectins and replicates, we further expressed the binding of each lectin as a ratio of ConA as an internal N-glycosylation control (Fig 2B). We found that all three IgG molecules showed strong ConA and LCA binding, indicating N-glycans with a high degree of core fucosylation. Conversely, IgG molecules were uniformly low in bisecting GlcNAc, β1,6 GlcNAc branching, and α2,6 sialylation, which is generally consistent with the glycan structure of murine IgG described in the literature as determined by the large number of studies on IgG glycosylation (reviewed in [1]).

**Figure 2.**
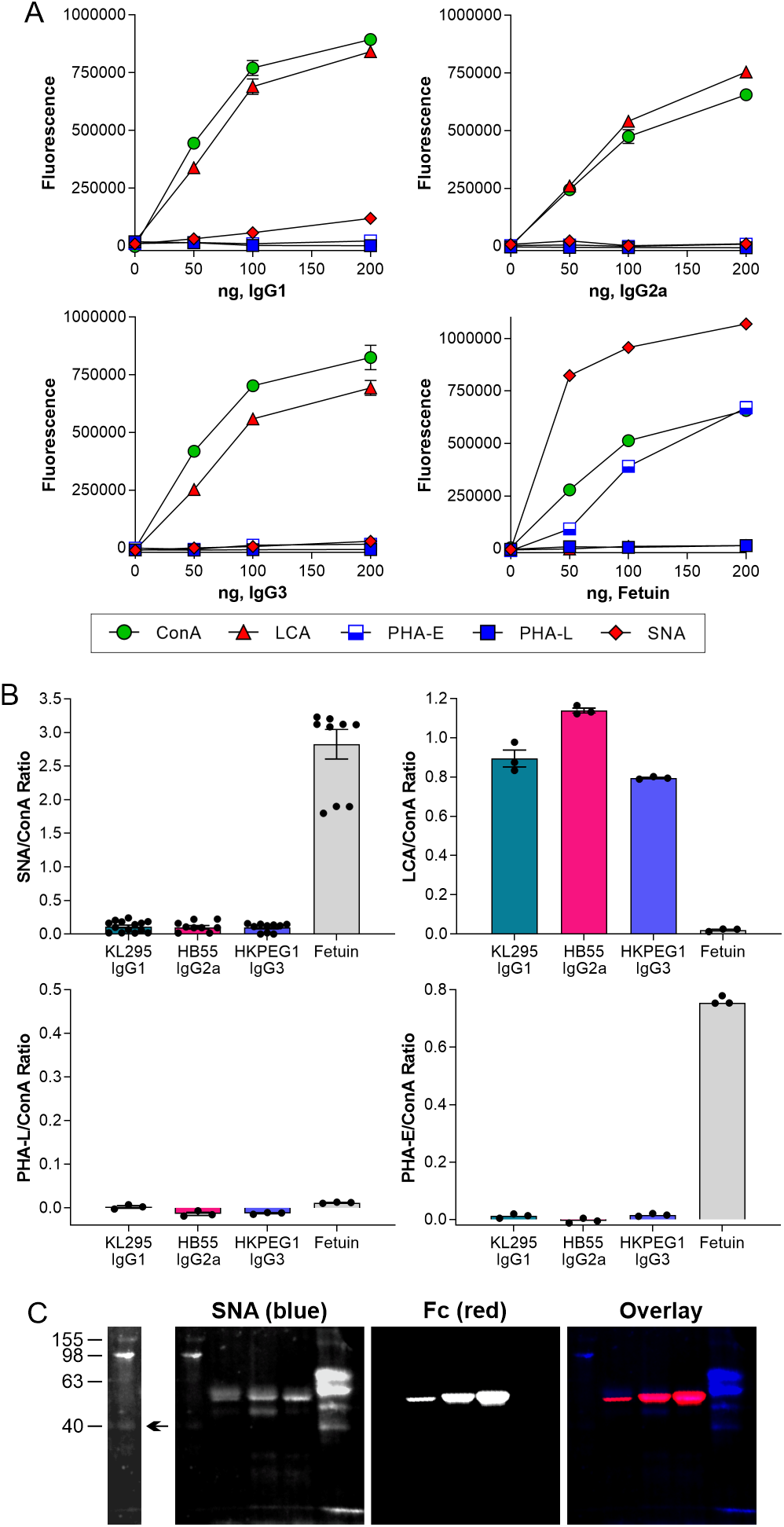
Hybridoma IgG have limited α2,6 sialylation and abundant core fucose. (A) ELISA of hybridoma IgG samples compared to bovine fetuin at a range of target protein concentrations and using a panel of lectins, including ConA (mannose), LCA (core fucose), PHA-E (bisecting GlcNAc), PHA-L (β1,6 branching), and SNA (α2,6 sialylation). N = 3. (B) Comparison of 100 ng each IgG subclass and fetuin based on the lectin:ConA ratio to normalize to total N-glycan. N = 3 for LCA, PHA-E, and PHA-L. N = 14 for IgG SNA and N = 9 for fetuin SNA. (C) Multiplexed Western blot of IgG subclasses using SNA and antimurine Fc antibody. All error bars show SEM.

Finally, we analyzed the IgG molecules by lectin-based Western blot. We confirmed a low but detectable degree of sialylation, as measured by SNA, that colocalized with the band that stained with anti-Fc antibody (Fig 2C).

In summary, we found that three murine hybridoma cell lines produce highly core fucosylated, poorly sialylated, and non-bisected biantennary IgG glycans.

### IgG Glycosylation Diverged from Cell Surface Glycosylation

Our data clearly showed that hybridoma IgG is poorly sialylated and highly core fucosylated. In an effort to determine why this pattern is seen, we first sought to determine whether these IgG molecules simply were not sialylated by the cell due to an unknown cellular mechanism, or if sialylation was low due to an IgG-intrinsic property. IgG was incubated with recombinant ST6Gal1, and the resulting IgG molecules were analyzed by lectin ELISA. We found that sialylation was significantly increased on all three IgG subclasses, ranging from a 75 % increase to over 200 % (Fig 3A), confirming both the presence of terminal galactose on hybridoma-produced IgG and its accessibility to the active site of ST6Gal1. This observation led us to hypothesize that the lack of sialylation must be due to a cell-intrinsic mechanism.

**Figure 3.**
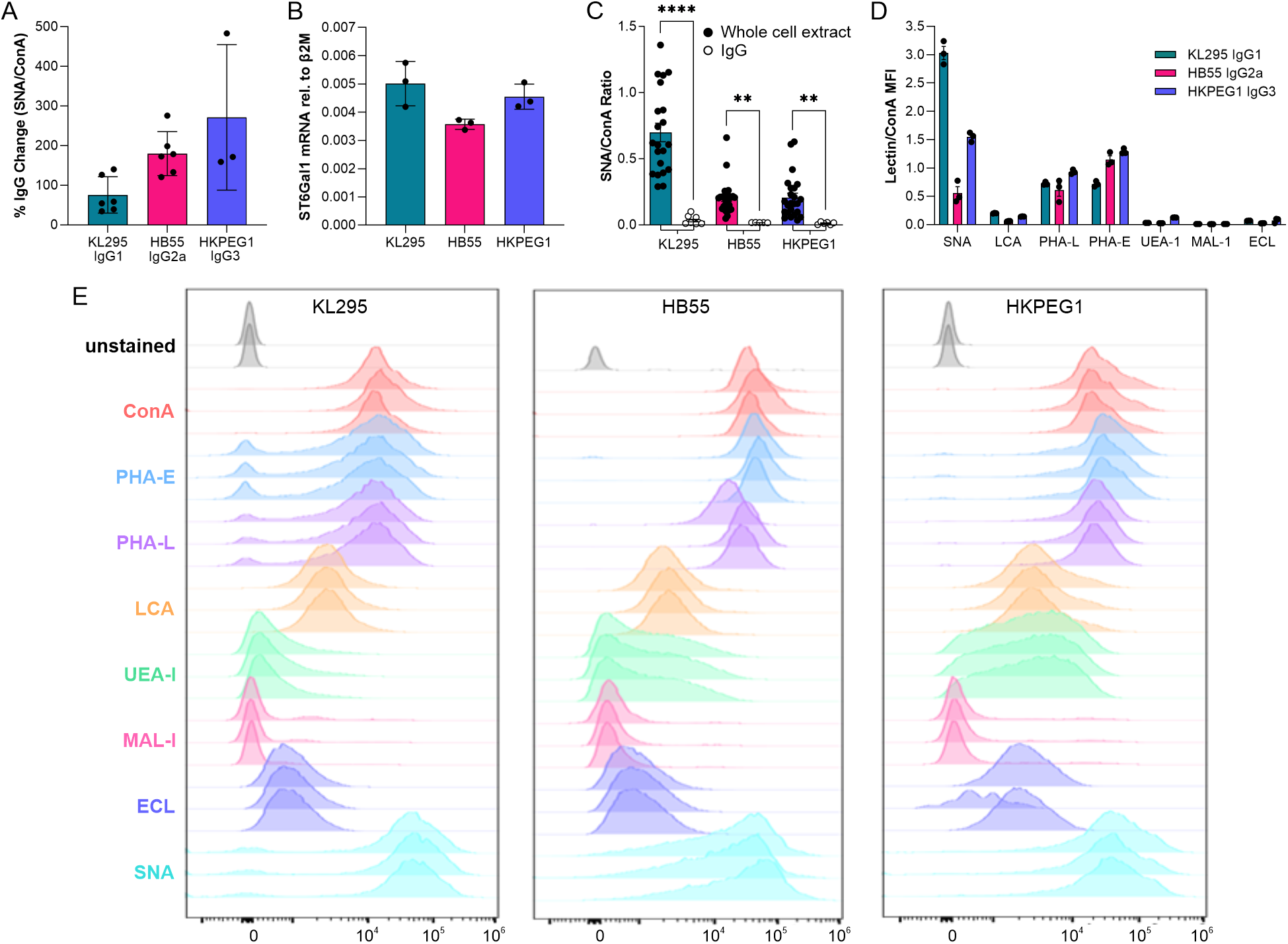
Hybridoma IgG glycans can be sialylated but differ from B cell surface glycans. (A) Hybridoma IgG subclasses were treated with recombinant ST6Gal1 and analyzed for the addition of sialic acid by SNA and ConA ELISA and expressed as a percent increase over the input IgG. N = 3-6. (B) Hybridoma ST6Gal1 mRNA level relative to β2 microglobulin transcript, as measured by quantitative PCR. N = 3. (C) SNA and ConA ELISA of whole cell extracts compared to IgG from each corresponding hybridoma cell line. N = 6 for each IgG; N = 21 for each whole cell extract. (D) Lectin:ConA ratios of the mean fluorescence intensities of each hybridoma cell line using a panel of lectins and flow cytometry. N = 3. (E) Raw flow cytometry histograms for each cell line and lectin used. N = 3. All error bars show SEM.

In order to determine why IgG is poorly sialylated in these cell lines, we next measured the expression of ST6Gal1 by quantitative PCR. We found strong expression of the gene at similar levels in the three lines (Fig 3B). We then harvested a whole cell extract from each line and compared the degree of sialylation to the purified IgG from the corresponding cell using lectin ELISA. Consistent with strong ST6Gal1 expression in each cell, we discovered a much higher degree of α2,6 sialylation in the total protein sample compared to isolated IgG (Fig 3C).

Finally, we used a selection of fluorescent lectins for flow cytometric analysis of live cells. Remarkably, we discovered that the cell surface glycans were highly divergent from those found on IgG (Figs 3D–3E). Plasma membrane glycans were highly α2,6 sialylated (SNA^high^), low in α1,6 core fucosylation (LCA^low^), and contained significantly higher bisecting GlcNAc (PHA-E) and branching beyond biantennary (PHA-L) compared to IgG from the same cells, suggesting differential pathways of glycan maturation in these hybridomas.

### Primary B cells Poorly Sialylate IgG

Since hybridomas are highly abnormal cells created by fusing B cells with a cancer cell line, we next sought to determine the degree to which an isolated but primary B cell can sialylate IgG in the absence of the circulatory environment. To accomplish this, we compared the relative degree of α2,6 sialylation of IgG isolated from murine plasma samples to IgG produced *ex vivo* by B cells from those same animals in a paired study. First, plasma was harvested and IgG purified. Second, CD19^+^ cells were isolated and analyzed by flow cytometry with SNA and cultured for 7 days with anti-CD40 and LPS stimulation to induce IgG production and secretion. Similar to previously published data [10], flow analysis showed high B cell SNA staining both before and after culture (Fig 4A), consistent with robust ST6Gal1 expression and activity. Comparisons of paired plasma and *ex vivo*-produced IgG using lectin ELISA revealed that plasma-localized IgG was significantly more sialylated than the IgG released in B cell culture. This was true when treating the data as replicates (Fig 4B) and as paired observations (Fig 4C). These findings suggest that B cells produce poorly sialylated IgG, just as we observed with hybridoma-produced IgG. Moreover, the data also lend additional support to the model proposed in our earlier work [10] in which a B cell-extrinsic mechanism drives sialylation *in vivo.*

**Figure 4.**
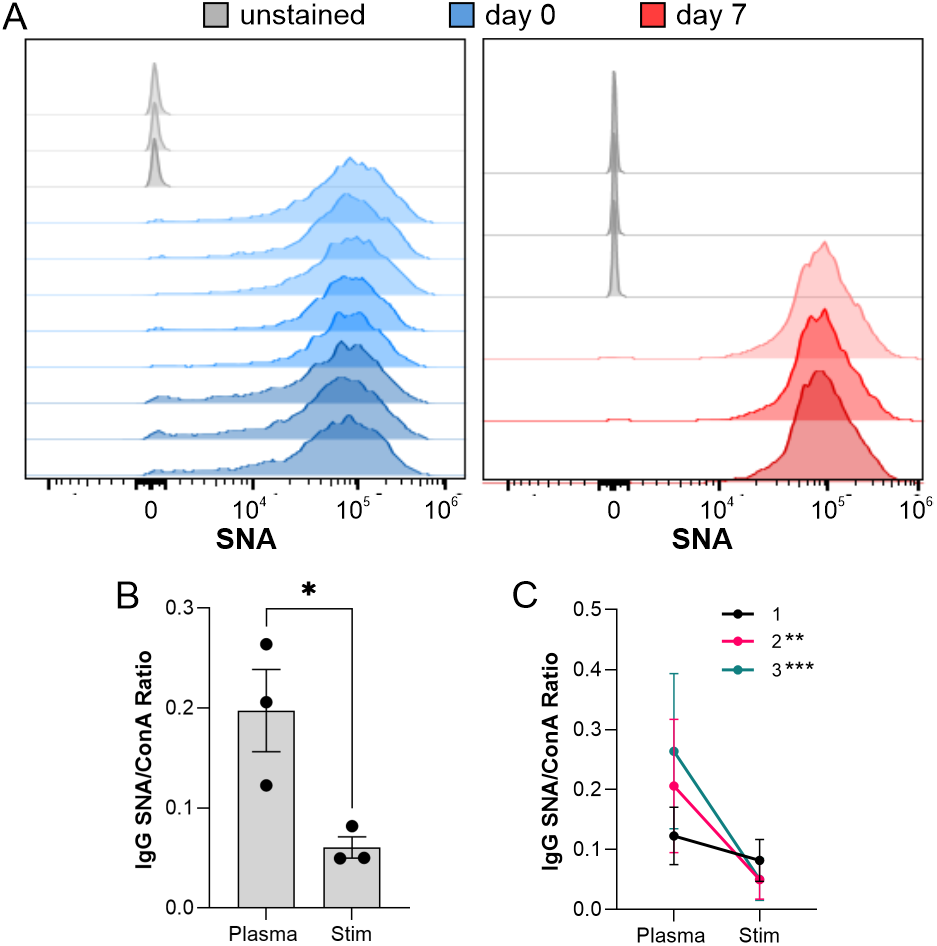
Primary murine B cells poorly sialylate IgG. (A) B cells isolated from wild type mice were analyzed by SNA flow cytometry before and after a 7 day stimulation to produce IgG. N = 3-8. (B) IgG purified from *ex vivo* B cell culture media and plasma from the same animal analyzed by SNA and ConA ELISA and graphed as a ratio. N = 3. (C) Alternative graph of the SNA:ConA ratios with connecting lines to indicate paired samples. All error bars show SEM. \*\*\**P*<0.001; \*\**P*<0.01.

### Intracellular IgG Trafficking Limits Sialylation

Our data established the existence of a unique glycosylation pattern present on IgG glycans compared to cell surface glycans in which IgG is characterized by low α2,6 sialylation and high α1,6 core fucosylation. We hypothesized that these contrasting glycosylation patterns may reflect differences in intracellular trafficking within the Golgi. In general, ST6Gal1 is thought to be localized to the *trans*-Golgi network, where it contributes to glycan maturation before secretion or localization to the plasma membrane [13, 14]. If IgG traverses the same pathway as other secretory and plasma membrane-localized glycoproteins, IgG sialylation should be similar to that of other glycoproteins at the cell surface, especially with the facility that recombinant ST6Gal1 appears to sialylate IgG *in vitro* (Fig 3A).

We performed confocal analyses to localize IgG and a variety of glycan structures using a panel of lectins in hybridomas (Fig 5A). Based on the Pearson’s Coefficient of colocalization in all three cell lines (Figs 5B–5D), we found that IgG robustly localized to intracellular compartments rich in N-glycans (ConA) and α1,6 core fucosylation (LCA), but poorly to regions rich in glycans containing β1,6 branched N-glycans (PHA-L), bisecting GlcNAc (PHA-E), and α2,6 sialylation (SNA). This is consistent with the glycosylation pattern on the purified IgG (Fig 2B), and highly divergent from the dominant glycoforms found on plasma membrane glycoproteins on the same cell (Figs 3C–3E). These data directly suggested that IgG comes into contact with Fut8 but has limited exposure to ST6Gal1 within the Golgi apparatus. In order to measure the relative intracellular IgG contact with these enzymes, we stained hybridomas for IgG and either Fut8 or ST6Gal1. We found that IgG colocalized to a much greater extent with Fut8 compared to ST6Gal1 (Fig 6A-C). These data collectively demonstrate that IgG was trafficked in such a way as to promote core fucosylation by Fut8 while limiting sialylation by ST6Gal1, thereby providing a mechanism underlying the relative inefficiency of B cell-mediated IgG sialylation.

**Figure 5.**
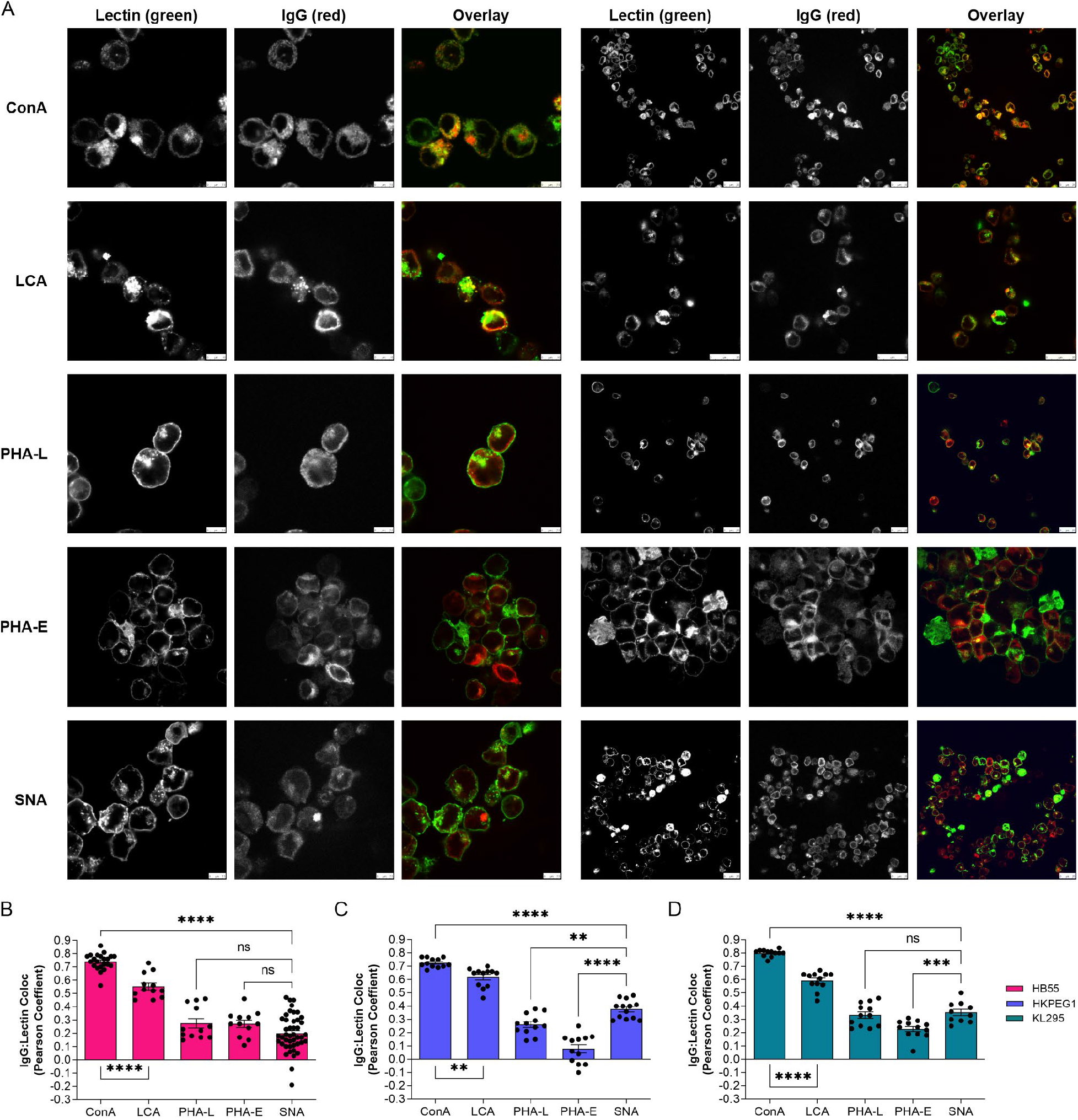
IgG poorly localizes to intracellular regions rich in α2,6 sialylation. (A) Hybridoma cells stained with a panel of lectins (green) and anti-IgG antibody (red) shown as individual channels and color overlay to illustrate colocalization (yellow). Two representative images for each lectin are shown. (B-D) Quantitation of colocalization between lectin-staining and IgG across many images for each hybridoma cell line, as indicated. N = 10-25. All error bars show SEM. \*\*\*\**P*<0.0001; \*\*\**P*<0.001; \*\**P*<0.01.

**Figure 6.**
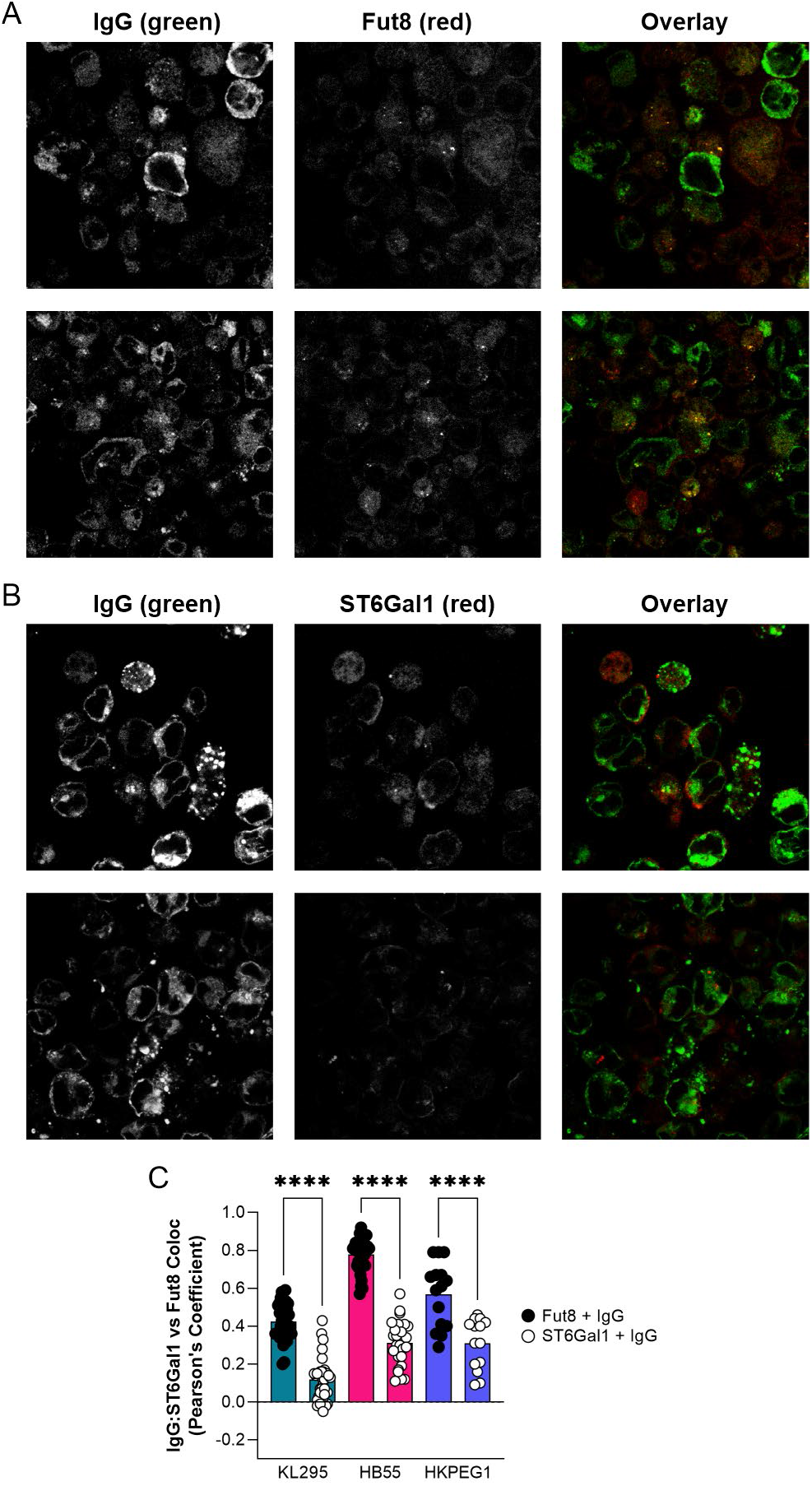
IgG colocalizes preferentially with Fut8 over ST6Gal1. (A) Representative images of hybridoma cells stained with anti-IgG (green) and anti-Fut8 (red) antibodies, shown as individual channels and color overlay to illustrate colocalization (yellow). (B) Representative images of hybridoma cells staining with anti-IgG (green) and anti-ST6Gal1 (red) antibodies, shown as individual channels and color overlay to illustrate colocalization (yellow). (C) Quantitation of colocalization between IgG and either Fut8 or ST6gal1 across many images for each hybridoma cell line. N = 12-30. All error bars show SEM. \*\*\*\**P*<0.0001.

## Discussion

IgG sialylation is recognized as a functional switch between pro-inflammatory and antiinflammatory activity, but the mechanism underlying the control of sialylation remains unknown. Previously, it has been assumed that secreted proteins are sialylated as they pass through the *transGolgi* network, and that sialylation levels are primarily regulated through up- or down-regulation of the enzyme ST6Gal1, and/or the degree to which terminal galactose residues, added by β1,4-galactosyltransferase 1 (B4GalT1), are available. Indeed, research has been published emphasizing a correlation between ST6Gal1 expression levels in B cells and the degree to which IgG carries sialic acids [15]. In addition, it has been known for decades that B4GalT1 is regulated in B cells as a function of rheumatic disease, RA in particular [1, 16, 17]. However, our previous work in generating a B cell-specific ST6Gal1 knockout showed that ST6Gal1 is dispensable in B cells, such that these mice express indistinguishable IgG sialylation profiles compared to wild type controls [10]. As a result, the present study was aimed at determining why B cells, which are strongly ST6Gal1 positive, do not appear to be centrally responsible for IgG glycan sialylation.

The paradox of distinct B cell and IgG glycosylation patterns was apparent in our 2016 study describing the B cell-specific ST6Gal1 knockout [10]. We showed that the knockout yielded B cells essentially void of α2,6 sialylation at the cell surface while IgG molecules in the circulation remained normally sialylated. This suggested that the mechanism by which B cell surface glycoproteins are sialylated is distinct from IgG sialylation. Here, we show detailed analysis of both B cell surface and IgG glycosylation in hybridoma cell lines and found remarkable differences. Specifically, the B cell surface was characterized by high α2,6 sialylation, moderate N-glycan bisection, and low core fucosylation, yet the IgG from those same cells showed very low to undetectable sialylation, no detectable bisection, and high core fucosylation. The low sialylation was not due to a lack of available terminal galactose residues, as treatment of these IgG molecules with recombinant ST6Gal1 *in vitro* yielded greatly increased sialylation. Moreover, we found poor sialylation of IgG produced by primary wild type B cells in culture despite their high degree of cell surface α2,6 sialylation, confirming that this phenomenon is not limited to a mutant mouse strain [10] and abnormal B cell hybridoma cell lines.

Confocal analyses of these cell lines then revealed a previously unrecognized mechanism to explain the inefficiency of B cells at IgG sialylation: vesicular trafficking. The trafficking of neoglycoproteins to Golgi compartments in which the various glycosyltransferases are found appears to diverge between the majority of plasma membrane-destined glycoproteins and IgG. Our data suggests that IgG is not robustly trafficked to the same location as ST6Gal1 in the *trons*-Golgi apparatus, but that it comes into contact with Fut8 consistently. Non-traditional protein secretion pathways in which the proteins avoid the Golgi apparatus are known to exist (reviewed in [18]), but it is believed that these pathways are primarily utilized in cell stress situations, and to our knowledge have not been previously implicated in IgG secretion. A pathway in which IgG avoids the Golgi altogether is also inconsistent with the abundant level of core fucosylation and galactosylation, as well as the lack of high mannose and hybrid structures. Instead, our data suggests that the secretory pathway may utilize variations in trafficking of individual glycoproteins within the *cis*-, *mediol-,* and *trons*-Golgi as a constant element of cell biology, rather than as a unique feature of cell stress. In such a model, glycoproteins may be shuttled in and out of the traditional pathway, allowing them to encounter selected enzymes but not others.

Selective trafficking is not conceptually novel. Proteins are directed to the various subcellular compartments through interactions with transmembrane-proximal domains and other signals, such as mannose-6-phosphate (M6P)[19]. Indeed, the existence of the N-terminal tag directing secreted proteins to the endoplasmic reticulum has been a staple of cellular biology since 1984 [20]. However, secretory IgG is unusual in that it lacks a transmembrane domain and the presence of M6P has never been reported on IgG glycans. If IgG is selectively trafficked through the Golgi as our data suggests, it would be predicted that a protein-intrinsic signal directing this pathway exists, although no evidence for such a protein has yet been described.

In summary, we have discovered that IgG is trafficked by antibody-producing cells such that sialylation is minimized and core fucosylation is maximized, thereby providing the body with a source of poorly sialylated IgG that can be dynamically sialylated elsewhere as a means of tightly and rapidly regulating IgG function. Thus, understanding the regulation of IgG glycosylation must take into consideration several aspects of cell biology, including (1) the expression and localization of glycosyltransferases, (2) the expression and localization of nucleotide-sugar transporter proteins, (3) the availability of nucleotide-sugar precursors, (4) the structure and steric availability on the target glycoprotein for enzymes to modify the glycans, and (5) the intracellular trafficking pattern of the specific glycoprotein. Further studies are needed to more clearly define the fine control and underlying mechanism behind IgG trafficking to more completely understand the factors that govern IgG functional control within B cells.

## Abbreviations Page

RA: rheumatoid arthritis
BcKO: B cell specific conditional knockout of ST6Gal1
HPAEC-PAD: high pH anion exchange chromatography with pulse amperometric detection
HILIC: hydrophilic interaction liquid chromatography
2-AB: 2-aminobenzamide

## Contributions

LMG and JYZ, data collection and analysis, manuscript writing and editing, experimental design; KMR, data collection and analysis; BAC, data collection and analysis, manuscript writing, experimental design, and funding.

## Acknowledgements

The authors wish to thank several former members of the Cobb laboratory for early contributions to this body of work; specifically, Edward Sim, Elianna Lai, and Kelsey Oliva.

